# Transmembrane protein KIRREL1 regulates Hippo signaling via a feedback loop and represents a potential therapeutic target in YAP/TAZ-active cancers

**DOI:** 10.1101/2022.02.28.482264

**Authors:** Yuan Gu, Yu Wang, Zhao Sha, Jian Li, Chenxi He, Fei Lan, Fa-Xing Yu

## Abstract

Dysregulation of the Hippo tumor suppressor pathway and hyperactivation of YAP/TAZ are frequently observed in human cancers and represent promising therapeutic targets. However, strategies targeting the mammalian Hippo pathway are limited due to the lack of a well-established cell surface regulator. By combining protein interactome data and clinical data, we have identified transmembrane protein KIRREL1 as an upstream regulator of the Hippo pathway. KIRREL1 interacts with Hippo pathway components SAV1 and LATS1/2 via its intracellular C-terminal domain and promotes LATS1/2 activation by MST1/2 (Hippo kinases), in turn inhibiting YAP/TAZ activity and target gene expression. Conversely, YAP/TAZ directly induce the expression of KIRREL1 in a TEAD1-4–dependent manner. In mouse liver tumors driven by YAP activation, KIRREL1 protein is robustly induced. Moreover, KIRREL1 expression positively correlates with canonical YAP/TAZ target gene expression in clinical tumor specimens and predicts poor prognosis. Finally, transgenic expression of KIRREL1 effectively blocked tumorigenesis in a mouse intrahepatic cholangiocarcinoma model, suggesting an important role of KIRREL1 in inhibiting cancer development. Together, these findings indicate that KIRREL1 constitutes a negative feedback mechanism regulating the Hippo pathway, and serves as a cell surface marker and potential drug target in cancers with YAP/TAZ dependency.

## Introduction

The Hippo pathway is a crucial regulator of organ size, tissue homeostasis, and tumorigenesis^1–5^. The activation of the Hippo pathway initiates a kinase cascade in which mammalian Ste20-like kinases 1/2 (MST1/2) and mitogen-activated protein kinase kinase kinase kinase 1-7 (MAP4K1-7) phosphorylate and activate large tumor suppressor 1/2 (LATS1/2) kinases, which in turn phosphorylate and inactivate Yes-associated protein (YAP) and WW domain-containing transcription regulator protein 1(WWTR1, also known as TAZ). Studies have identified several proteins, including moesin-ezrin-radixin like tumor suppressor (NF2, also known as Merlin), salvador homolog 1 (SAV1), and WW and C2 domaincontaining proteins (WWC1/2/3), that play important roles in mediating Hippo signaling. Moreover, diverse upstream signals, such as cell-cell contact, cell polarity, mechanical cues, cellular energy status, and G-protein-coupled receptor (GPCR) ligands have been shown to regulate the activity of the Hippo pathway^3,6–10^. Downstream of Hippo signaling, YAP/TAZ interact with TEA domain transcription factors 1-4 (TEAD1-4) to modulate the expression of genes involved in cell proliferation, differentiation, and death, in turn affecting organ growth and tumorigenesis.

Dysregulation of Hippo signaling and resulting YAP/TAZ activation play key contributions to the development of human cancers. Indeed, YAP/TAZ activation is frequently observed in various cancer types including liver, stomach, breast, lung, and colon^11–16^. Mechanistically, YAP/TAZ activity has been shown to induce cancer stem cell traits, proliferation, metastasis, and chemoresistance^15^. In genetic mouse models, activation of YAP/TAZ also leads to tumorigenesis^6,17^, suggesting YAP/TAZ inactivation as a promising strategy to inhibit cancer development. Despite current efforts to target Hippo signaling in cancer^18^, inhibition of YAP/TAZ activity remains challenging because most signaling nodes in the pathway are not druggable, and attempts to inhibit YAP/TAZ are mostly focused on disrupting YAP/TAZ–TEAD1-4 interaction ^19,20^. Therefore, the identification of new, druggable regulators of the Hippo pathway is an urgent need.

Cancer cells can specifically turn on expression of certain cell surface proteins that can then be used as biomarkers for cancer diagnosis ^21^. These cancer cell-associated cell surface proteins can serve as excellent therapeutic targets as they are easily accessible by large antibody- and cell-based therapies ^21^. The identification of a novel cell surface regulator for the mammalian Hippo pathway that can be can be targeted to suppress YAP/TAZ activation can greatly advance the development of therapies targeting the Hippo pathway.

In this study, we have identified the cell surface protein KIRREL1 as a Hippo pathway component inhibiting YAP/TAZ activity. Mechanistically, the intracellular C-terminal domain of KIRREL1 interacts with both LATS1/2 and SAV1, and the latter brings in MST1/2 to phosphorylate and activate LATS1/2. We demonstrate that *KIRREL1* is a direct target gene of YAP/TAZ, and high KIRREL1 expression predicts poor prognosis in gastric cancer. Finally, in a mouse cholangiocarcinoma model, we show that transgenic expression of KIRREL1 effectively blocked tumorigenesis. Thus, KIRREL1 serves as a negative regulator of YAP/TAZ, a biomarker in YAP/TAZ-active cancers, and potentially a therapeutic target in YAP/TAZ-driven tumors.

## Results

### Identification of KIRREL1 as a negative YAP regulator

The use of next-generation sequencing in the past decade has generated a vast amount of transcriptomic data for clinical tumor samples. As the activity of the Hippo pathway is faithfully reflected by the expression of YAP/TAZ target genes, we focused on two known YAP/TAZ target genes and well-established reporters of YAP/TAZ activity, *CTGF* and *CYR61.* We compared the prognostic significance of the Hippo pathway in different cancers by calculating the hazard ratio (HR) of *CTGF* and *CYR61* levels in different TCGA cancer datasets. Notably, both *CTGF* and *CYR61* levels showed the most significant prognostic value in stomach adenocarcinoma (STAD) (Fig. S1a-d, Table S3). Enrichment analysis combining *CTGF* and *CYR61* levels was used to define the on/off/moderate status of Hippo and YAP in stomach cancer patients. Prognostic analysis showed that stomach cancer patients with Hippo off/YAP on status had a significantly worse prognosis (Fig. S1e-f, Table S4). Adopting the stomach cancer clinical data, we screened for clinically significant Hippo pathway regulators. Among the 21,399 genes analyzed, 3,646 genes showed significant prognostic value and correlated with *CTGF* and *CYR61* expression levels (Fig. 1A, Table S2).

**Fig. 1.**
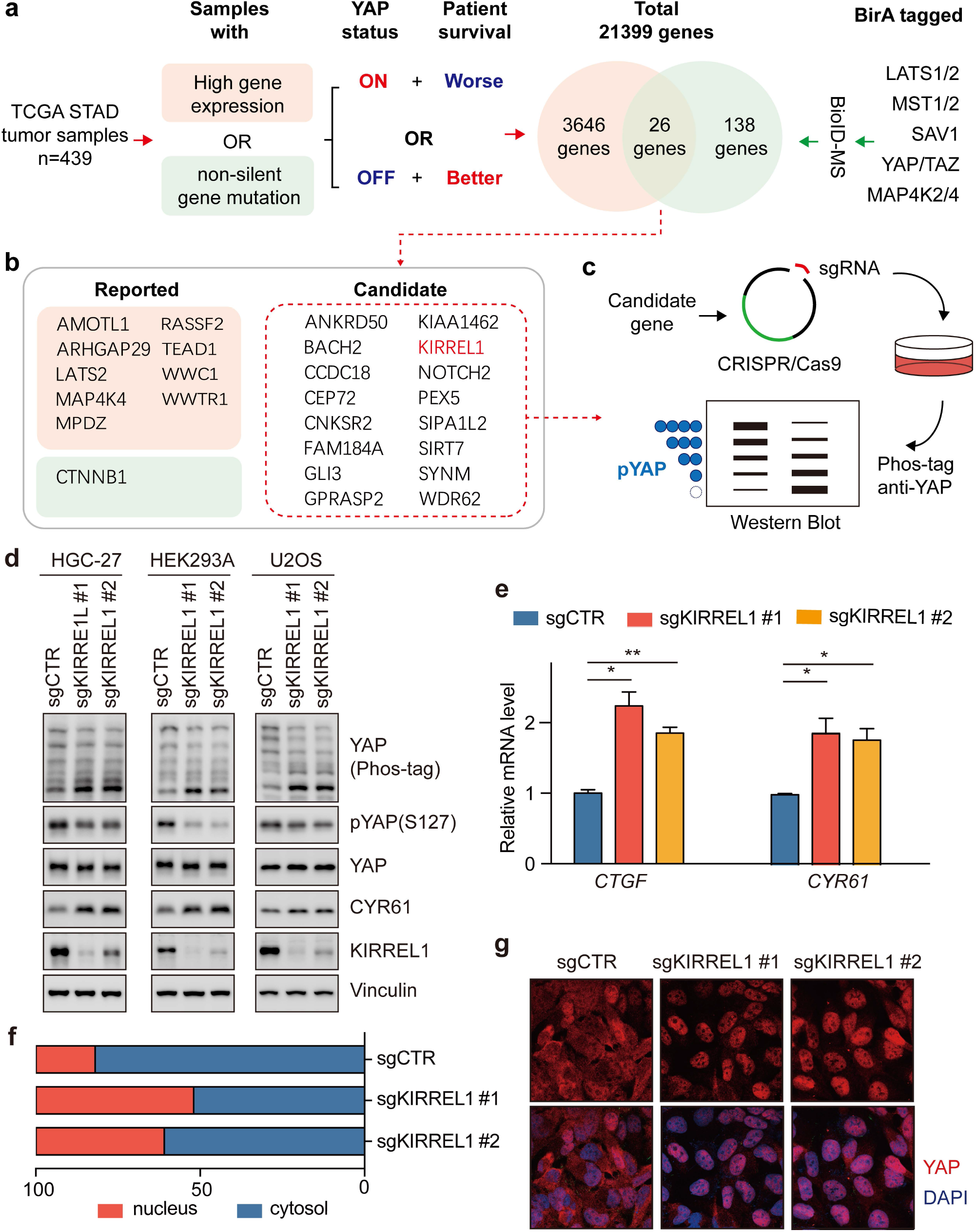
Identification of KIRREL1 as a negative regulator of YAP. (A) Scheme for the primary screen, which integrated TCGA stomach cancer clinical data and LATS1/2, MST1/2, SAV1. YAP/TAZ, and MAP4K2/4 interactome data as detected by BioID-MS. (B) List of 26 candidate genes, among which 10 were reported as known Hippo pathway regulators and other 16 genes entered secondary genetic screen. (C) Scheme for the secondary screen. (D) HGC-27, HEK293A, and U2OS KIRREL1-knockout cells showed de-phosphorylation of YAP. Phos-tag SDS-PAGE was performed to separate phosphorylated and non-phosphorylated YAP protein by mixing Phos-tag™ acrylamide with acrylamide solution to allow for polymerization to occur. The relevance of YAP phosphorylation level and Phos-tag blotting signals was indicated in Fig. 1C. (E) qPCR showed KIRREL1 KO increased *CTGF* and *CYR61* levels in HEK293A cells. Mean and standard error were presented (*p<0.05, **p < 0.01, ***p < 0.001, ****p <0.0001, ns = not significant; n=2; t test). (F) Immunofluorescence showed increased nuclear YAP localization in KIRREL1 KO cells. Cells with more cytoplasmic or nuclear YAP staining were artificially counted. (G) Representative immunofluorescence images of Fig. 1F.

Next, using BioID proximity labeling coupled with mass spectrometry, we searched for proteins interacting with known Hippo pathway components including LATS1, LATS2, MST1, MST2, SAV1, YAP, TAZ, MAP4K2, and MAP4K4. In total, 138 proteins were identified as candidates interacting with at least one Hippo pathway component (Figure 1a, Table S1). We then compared our interactome data with the candidates identified in the clinical screen and identified 26 shared candidate genes, among which 16 genes had not been reported as Hippo regulators (Fig. 1b).

In a secondary genetic screen, we employed CRISPR/cas9 technology to knock out candidate genes one by one in HEK293A cells and HGC-27 gastric cancer cells, and evaluated their effects on YAP phosphorylation using Phos-tag electrophoresis (Fig. 1c). Among 16 candidate genes, only *KIRREL1* (Kin of IRRE-like protein 1) knockout cells showed a robust reduction in YAP phosphorylation in both cell lines, and the effect of KIRREL1 was comparable to that of NF2, which was used as a positive control (Fig. S1g). The effect of KIRREL1 on YAP phosphorylation was further validated using independent sgRNA in three different cell lines (Fig. 1d). Dephosphorylated YAP translocates into the nucleus to activate gene expression ^17,22^. Consistently, in *KIRREL1*-deficient cells, increased nuclear YAP was detected, as shown by immunofluorescence staining (Fig. 1f, g), and *CTGF* and *CYR61* levels were also significantly induced (Fig. 1d, g). These results suggest that KIRREL1 is a negative regulator of YAP.

### Intracellular C-terminal domain of KIRREL1 is required for YAP inhibition

KIRREL1, also known as KIRREL, NEPH1, or NPHS23, is a member of the nephrin-like protein family and is involved in normal development and maintenance of glomerular permeability, as *Kirrel1* knockout mice develop heavy proteinuria and early death^23^. KIRREL1 is a transmembrane protein containing five immunoglobulin-like domains in its extracellular domain (ECD), a transmembrane domain (TM), and an intracellular domain (ICD) (Fig. 2a). Four major isoforms of KIRREL1 have been annotated, all of which share common TM domain and ICD (Fig. 2a). When ectopically expressed in wild-type or *KIRREL1* knockout HEK293A cells, KIRREL1 isoform 2 showed the strongest induction of YAP phosphorylation (Fig. 2b, S2a). Subsequently, we used isoform 2 as full-length KIRREL1 in most experiments in this paper unless otherwise indicated. KIRREL1 overexpression significantly increased YAP phosphorylation and decreased the expression of YAP target genes in different cancer cell lines, including gastric (HGC27 and AZ521), liver (HepG2), and osteosarcoma (U2OS) (Fig. 2c, 2d). Moreover, KIRREL1 overexpression induced cytoplasmic localization of YAP in multiple cell lines (Fig. 2e-f). Notably, stomach cancer samples from the TCGA database showed increased levels of all KIRREL1 isoforms, suggesting tumor suppressive roles of KIRREL1 (Fig. S2b).

**Fig. 2.**
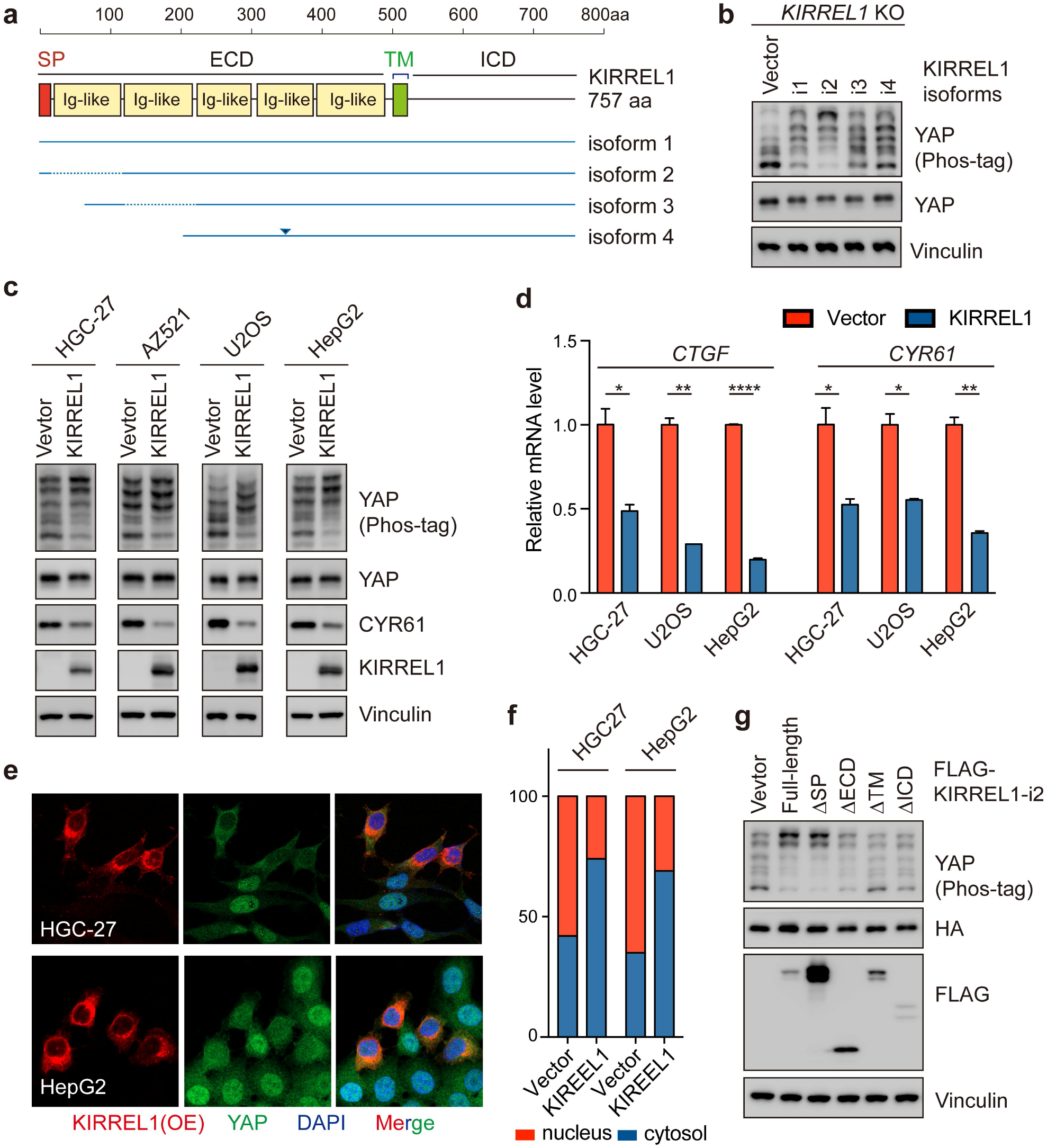
Intracellular C-terminal domain of KIRREL1 is required for YAP inhibition. (A) Domains and isoforms of KIRREL. SP, signal peptide. ECD, extracellular domain. TM, transmembrane domain. ICD, intracellular domain. (B) All isoforms of KIRREL1 increased YAP phosphorylation level while isoform 2 showed the strongest effect in KIRREL1 KO HEK293A cells. Isoform 2 was used as fulllength KIRREL1 in subsequent experiments except for extra statements. Since KIRREL1 had a signal peptide in its N terminal region and functional domain in its C terminal region, we preferred to overexpress no-tag KIRREL1 in most experiments as it mimicked endogenous KIRREL1 more. (C) KIRREL1 overexpression increased YAP phosphorylation in HGC-27, AZ521, U2OS, and HepG2 cells. (D) qPCR showed that KIRREL1 decreased *CTGF* and *CYR61* mRNA levels in HGC-27, U2OS, and HepG2 cells. Mean and standard error were presented (*p<0.05, **p < 0.01, ***p < 0.001, ****p <0.0001, ns = not significant; n=2; t test). (E) Immunofluorescence showed increased cytosolic localization of YAP in KIRREL1-overexpressing cells. (F) Quantitative result of Fig. 1e. Cells with higher cytoplasmic or nuclear YAP staining were artificially counted. (G) Overexpression of KIRREL1 mutants with deletions of different domains in KIRREL1 KO HEK293A cells showed that the regulation of the Hippo pathway by KIRREL1 was dependent on the transmembrane (TM) domain and cytoplasmic (C) domain of KIRREL1 and was partly impaired by deletion of its extracellular part.

To determine which amino acid (aa) motifs in KIRREL1 are critical for Hippo pathway regulation, we performed a series of amino acid truncations and analyzed their effects on YAP phosphorylation. As shown in Figure 2g, the TM domain and ICD of KIRREL1 were indispensable, while the ECD was partially required for YAP inhibition. A more detailed mapping showed that regions aa401-500 and C-terminal 57aa (Fig. S2c, d) played important roles in regulating YAP phosphorylation. We noticed that aa401-500 spanned the TM domain, indicating that plasma localization of KIRREL1 was required for YAP inhibition (Fig. S2c, also see below). We also examined the C-terminal 57aa region, which might potentially be involved in the crosstalk with Hippo pathway components. Within this 57aa region, however, 20aa deletion mutants did not fully impair KIRREL1’s function (Fig. S2e). We next checked tyrosine residues (Y) 605 and 606, which have been reported to be phosphorylated by tyrosine-protein kinase Fyn^24^. Mutations in these two residues to alanine (A), aspartic acid (D), or glutamic acid (E) did not impair the function of KIRREL1 (Fig. S2f). Finally, we examined the effect of serine 573 to leucine mutation (S573L), which has been reported to be frequently mutated in European populations^25^. As shown in Fig. S2g, S573L mutation induced YAP phosphorylation in a manner comparable to wildtype KIRREL1 (Fig. S2g), indicating that S573 is dispensable for YAP regulation. Together, these results demonstrate a critical role of the C-terminal 57aa within the ICD domain, probably in a threedimensional fold, in regulating Hippo pathway activity.

### KIRREL1 activates LATS1/2 by interacting with LATS1/2 and SAV1 through ICD

To investigate how KIRREL1 regulates YAP activity, we first tested if this regulation was dependent on Hippo kinase signaling. As shown by kinase assay, KIRREL1 expression effectively increased the phosphorylation and kinase activity of LATS1 (Fig. 3a). We then overexpressed KIRREL1 in a series of cell lines with knockout of Hippo pathway components and found that the regulation of YAP activity by KIRREL1 was dependent on LATS1/2, SAV1, MST1/2, NF2 and MOB1A/B, but not on MAP4Ks (MAP4K3-7), WWC1/2/3, and Motins (AMOT, AMOTL1, and AMOTL2). We then analyzed if KIRREL1 interacted with different Hippo pathway components. Our BioID-MS data showed that KIRREL1 interacted with LATS1/2 or SAV1 (Table S1), which was further supported by reciprocal co-immunoprecipitation assays using FLAG-tagged KIRREL1 expressed in different knockout cell lines (Fig. 3c-e). KIRREL1 also indirectly interacted with MST1/2, likely via SAV1 as the interaction between KIRREL1 and MST1/2 was significantly reduced in *SAV1* knockout cells, and strengthened in SAV1-overexpressing cells (Fig. 3f). These data demonstrate an extensive crosstalk between KIRREL1 and Hippo signaling components.

**Fig. 3.**
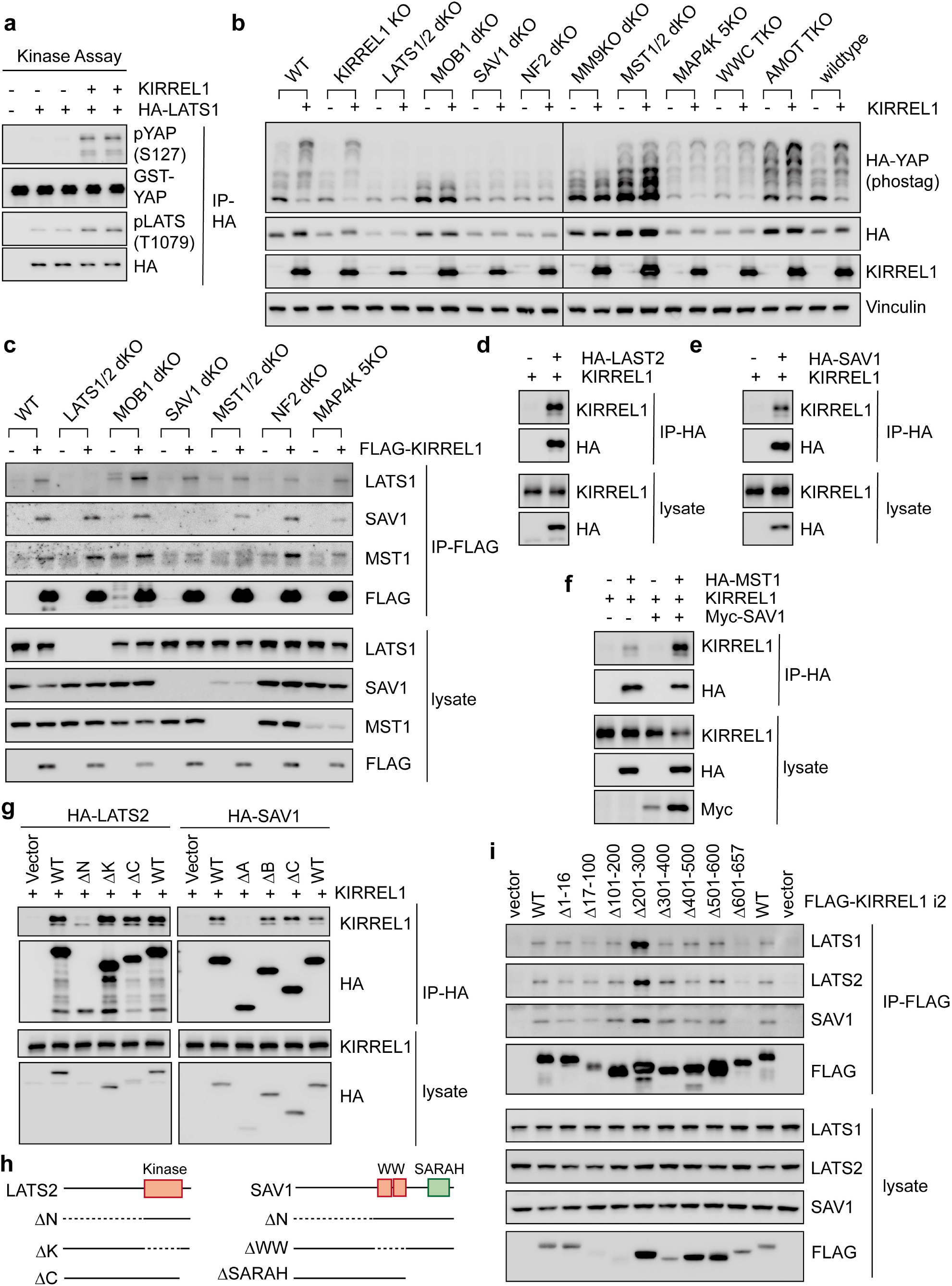
KIRREL1 activates LATS1/2 by interacting with LATS1/2 and SAV1 through ICD. (A) Kinase assay showed that KIRREL1 increased kinase activity of LATS1. HA-tagged LATS1 was immunoprecipitated from HEK293A cell lysates and its activity was measured in an *in vitro* kinase assay with purified YAP protein as substrate. (B) Overexpression of KIRREL1 in different Hippo pathway component KO cells showed its dependency on other Hippo regulators. (C) Co-immunoprecipitation of FLAG-tagged KIRREL1 in different Hippo component KO cells detected interaction of KIRREL1 with LATS1, SAV1, and MST1, whereas no interaction between KIRREL1 and MST1 was observed in SAV1 KO cells. (D) Co-immunoprecipitation assay detected the interaction between HA-tagged LATS2 and no-tag, overexpressed KIRREL1. (E) Co-immunoprecipitation assay detected the interaction between HA-tagged SAV1 and no-tag, overexpressed KIRREL1. (F) Co-immunoprecipitation assay showed that co-expression of SAV1 boosted the interaction between KIRREL1 and MST1. (G) Left: Full-length, 668-973 aa(ΔB), 974-1052 aa(ΔC) but not 1-667aa deleted (ΔA) LATS2 mutants interacted with KIRREL1. Right: Full-length, 199-267 aa deleted (ΔB), 268-383 aa deleted (ΔC) but not 1-198 aa deleted (ΔA) SAV1 mutants interacted with KIRREL1. (H) Structure of LATS1 and SAV1 mutants. (I) Co-immunoprecipitation of FLAG-tagged KIRREL1 100 aa deletion mutants in HEK293A showed that interaction of KIRREL1 with LATS1, LATS2, and SAV1 was dependent on its C-terminal 57aa.

We further mapped the protein domains responsible for the interaction between KIRREL1 and LATS1/2 or SAV1. The kinase domain of LATS2 is located at aa668-973 (Fig. 3g). We have generated LATS2 mutants with N-terminal domain (aa1-667, ΔN), kinase domain (aa668-973, ΔK), or C-terminal domain (aa974-1052, ΔC) deletion. We found that the N-terminal region of LATS2 was responsible for LATS2 interaction with KIRREL1 (Fig. 3g). To determine whether the two WW domains and SARAH domain of SAV1 mediate SAV1 interaction with KIRREL1, we generated SAV1 mutants with N-terminal domain (aa1-198, ΔN), WW domain (aa199-267, ΔWW), or SARAH domain (11268-383, ΔSARAH) deletion. We found that the N-terminal region of SAV1 was responsible for SAV1 interaction with KIRREL1 (Fig.3h). SAV1 and MST1/2 are known to interact via their respective SARAH domains^26^. Thus, SAV1 might work as an adaptor between KIRREL1 and MST1 and promote KIRREL1-MST1 interaction (Fig. 3c, f). As for the different KIRREL1 deletion mutants, the removal of C-terminal 57aa impaired the interaction between KIRREL1 and LATS1/2 or SAV1 (Fig. 3i), which might explain the inability of this mutant to induce YAP phosphorylation (Fig. S2d). Collectively, these results suggest that KIRREL1 inhibits YAP by inducing LATS1/2 kinase activity via interaction with LATS1/2 and SAV1, and SAV1 likely recruits MST1/2 to phosphorylate and activate LATS1/2.

### Tight junction localization of KIRREL1 is essential for integrating Hippo signaling

KIRREL1 aa401-500 deletion mutant failed to induce YAP phosphorylation, suggesting that the transmembrane domain is essential for the function of KIRREL1 (Fig.2d). Indeed, a refined mapping indicated that aa401-420, which corresponds to the exact TM domain of KIRREL1, is required for KIRREL1 to inhibit YAP (Fig. 4a), suggesting that plasma membrane localization of KIRREL1 plays an important role in KIRREL1 function. Immunofluorescence staining showed that both endogenous and ectopically expressed KIRREL1 co-localized with tight junction protein ZO-1 (Fig. 4b, c). However, KIRREL1 ΔTM (aa401-420) mutant was not targeted to tight junction (Fig.4b). While there is a potential membrane sorting signal peptide (aa1-16) at the extreme N-terminal of KIRREL1, deletion of this signal peptide did not affect KIRREL1 membrane localization and YAP phosphorylation (Fig. S3a,b). It has been shown previously that simultaneous recruitment of LATS1/2 and MST1/2 by NF2 and SAV1, respectively, to the plasma membrane is critical for LATS1/2 activation by Hippo kinases^27^. Given that KIRREL1 interacts with SAV1, KIRREL1 may provide an anchor for the SAV1-MST1/2 complex at the plasma membrane. On the other hand, in *NF2* knockout cells, although the effect of KIRREL1 on YAP phosphorylation was largely abolished, the interaction between KIRREL1 and LATS1/2 are largely normal (Fig. 3b, c). We further validated the requirement for NF2 in KIRREL1-induced phosphorylation of YAP in two independent *NF2* knockout cells (Fig. 4d). We have previously shown that NF2 is involved in the establishment of tight junctions^28^. In *NF2-* deficient cells, the organization of tight junctions was significantly disrupted, which consequently reduced the tight junctional localization of KIRREL1 (Fig. 4c). Therefore, we propose that KIRREL1 may recruit both SAV1-MST1/2 and LATS1/2 to tight junctions to facilitate LATS1/2 activation by MST1/2, whereas NF2 may indirectly affect this process by regulating the integrity of tight junctions (Fig. 4e).

**Fig. 4.**
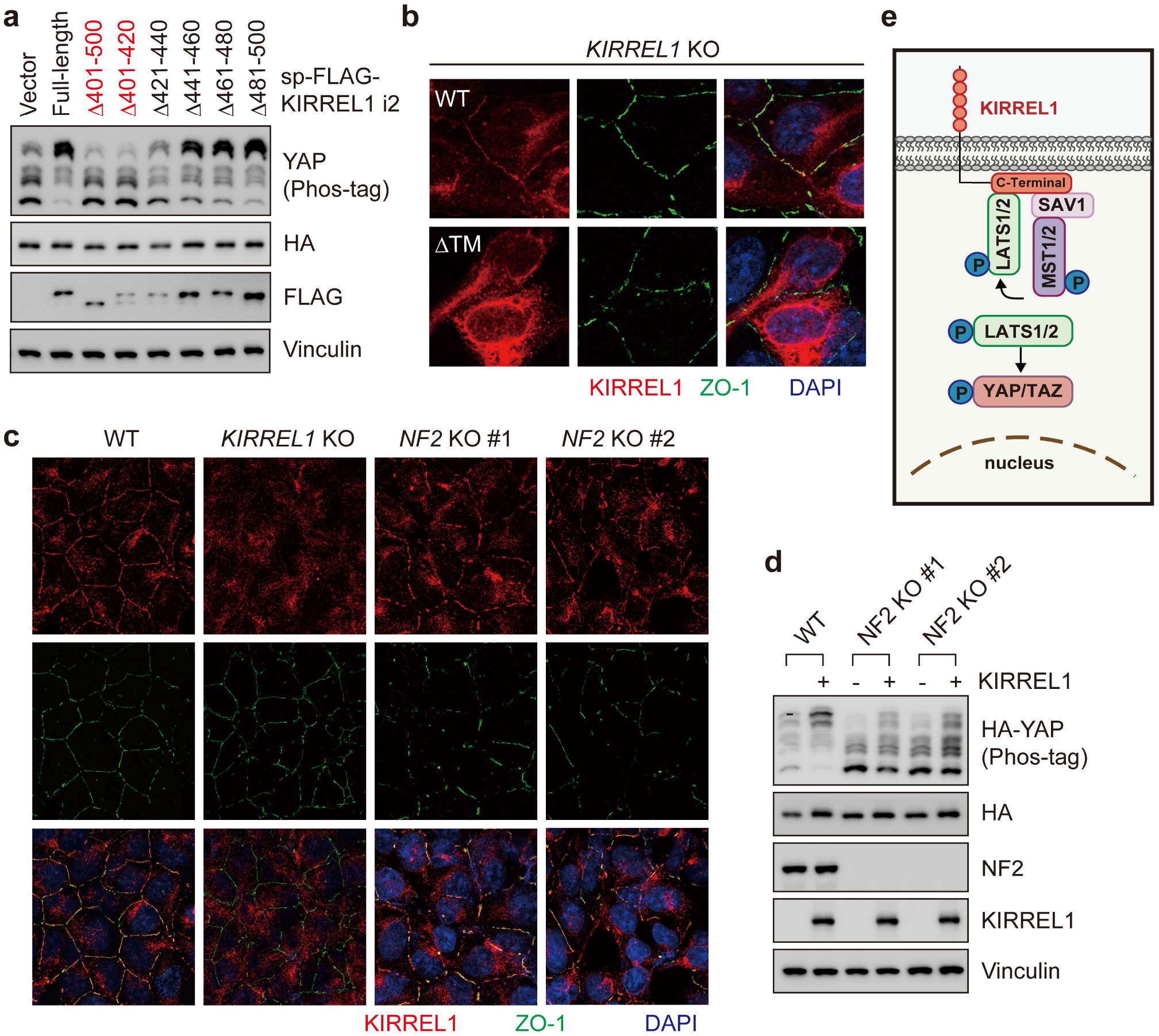
Tight junction localization of KIRREL1 is essential for integrating Hippo signaling. (A) Overexpression of KIRREL mutants each deleting 20aa within 401-500aa in KIRREL KO HEK293A cells showed that the effect of KIRREL on YAP phosphorylation was dependent on 401-420aa TM domain. (B) Immunofluorescence staining showed that full-length KIRREL1 co-localized with ZO-1 at tight junctions, while TM-domain-deleted mutant lost tight junction co-localization. Full-length KIRREL1 and TM-domain-deleted mutant were expressed in KIRREL1 KO HEK293A cells. (C) Immunofluorescence staining showed that NF2 KO largely impaired tight junction, marked by ZO-1 together with KIRREL1 co-localization. Notably, cytoplasmic signals of KIRREL1 antibody in immunofluorescence kept in KIRREL1 KO cells indicated it was non-specific signals. (D) The effect of KIRREL1 on YAP phosphorylation was largely impaired in NF2 KO cells. (E) Working model of the regulation of YAP activity in tight junction by KIRREL1.

### KIRREL1 partially mediates Hippo signaling activation by high cell density

The localization of KIRREL1 at tight junctions and its effect on YAP phosphorylation suggest a role of KIRREL1 in mediating cell-cell contact signal to the Hippo pathway, as cell-cell contact or high cell density is an important upstream signal regulating the Hippo pathway^22^. On the other hand, YAP phosphorylation is also robustly regulated by serum factors in cell media^10^. The number (or density) of cells in a culture dish is a determinant of the turnover rate of growth factors in serum. Thus, serum availability is a confounding variable when interpretating cell density data. We have shown previously that, among different cell lines tested, U2OS was least sensitive to serum^10^. Hence, U2OS cells were used to study the role of KIRREL1 in crowding-induced YAP phosphorylation.

We have monitored the formation of tight junctions and KIRREL1 localization in U2OS from single cells to a confluent culture. We noticed that KIRREL1 was diffused in single cells and localized to tight junctions when the latter was established upon cell-cell contact (Fig. 5a). On the other hand, deletion of *KIRREL1* in U2OS cells had no significant effect on tight junctions (Fig. 5a). The response of YAP to cell density in U2OS cells was robust, and YAP phosphorylation gradually increased when cells were cultured to higher cell densities (Fig. 5b). Interestingly, we found that the response of YAP phosphorylation towards higher cell density was significantly impaired in *KIRREL1* knockout cells (Fig.5b). Moreover, the inhibition of canonical YAP target gene expression—including *CYR61, ANKRD1, and AMOTL2—*in *KIRREL1* knockout cells followed a much slower kinetics (Fig. 5c-e). These results, together with the requirement of NF2-regulated tight junction formation (Fig. 4c), indicate that KIRREL1 plays a role, at least partially, in transducing the signals of cell-cell contact to Hippo kinase signaling.

**Fig 5.**
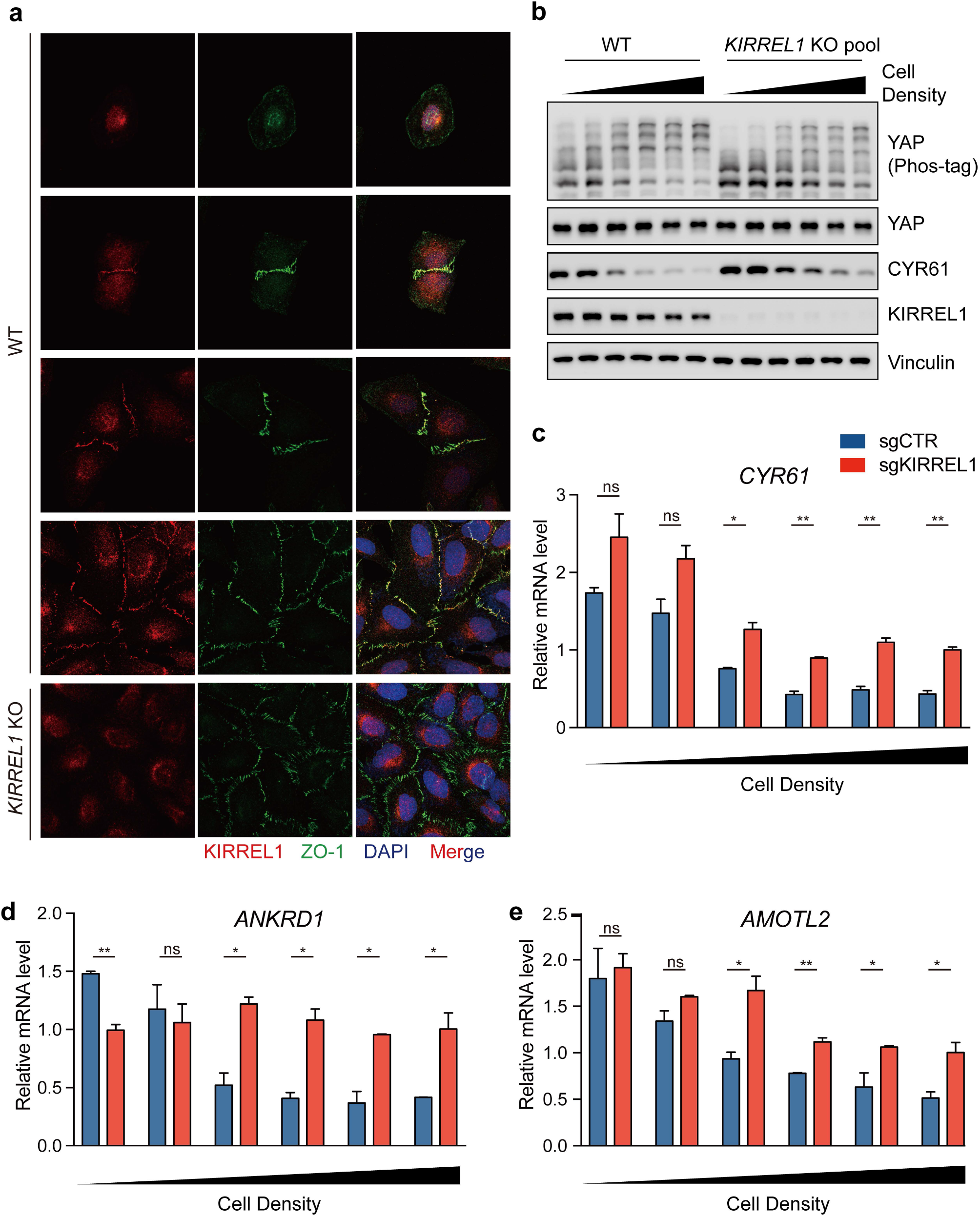
KIRREL1 partially mediates Hippo signaling activation by high cell density. (A) Immunofluorescence staining showed the localization of KIRREL1 under different cell densities. Higher cell density led to the formation of tight junctions marked by ZO-1 signals. (B) YAP phosphorylation level increased with higher cell density, while KIRREL1 KO significantly blunted this response. U2OS cells were seeded into 24-well, 12-well, 6-well plant, and 6 cm dish for different cell densities. After attachment, cells were harvested for immunoblotting. (C) qPCR showed that *CYR61* mRNA level decreased with higher cell density, while KIRREL1 KO significantly blunted this response. (D) qPCR showed that YAP target gene *ANKRD1* mRNA level decreased with higher cell density, while KIRREL1 KO significantly blunted this response. (E) qPCR showed that target gene *AMOTL2* level decreased with higher cell density, while KIRREL1 KO significantly blunted this response. Mean and standard error were presented (*p<0.05, **p < 0.01, ***p < 0.001, ****p <0.0001, ns = not significant; n=2; t test).

### *KIRREL1* is a YAP/TAZ target gene

KIRREL1 was first identified as a potential Hippo pathway component by analysis of stomach (STAD) transcriptome data, in which *KIRREL1* was highly expressed in the Hippo off/YAP on group (Fig. 6a). The expression of *KIRREL1* in STAD highly correlated with *CTGF* and *CYR61* levels (Fig. 6b). We therefore tested whether KIRREL1 could be a YAP target gene. We found that overexpression of YAP 2SA, an active YAP mutant^29^, in HGC-27 cells increased the mRNA and protein levels of KIRREL1 (Fig. 6c). On the contrary, overexpression of *LATS2* or knockout of *YAP/TAZ* showed an opposite effect (Fig. 6d, e). Moreover, when different cells were cultured at high cell densities to turn on Hippo signaling, KIRREL1 protein levels were significantly reduced, although not as dramatic as CYR61 (Fig 6f). We also examined the effect of YAP activation on KIRREL1 expression *in vivo.* Overexpression of constitutively active YAP 5SA in mouse livers by hydrodynamic injection induced hepatocellular carcinomas^30^ (Fig. 6g). Notably, the expression of KIRREL1 in these YAP-driven liver tumors was significantly higher than in normal adjacent tissues (Fig. 6h), indicating that YAP regulates the expression of KIRREL1. Moreover, as indicated by ENCODE chromatin immunoprecipitation sequencing (ChIP-Seq) data, different TEAD proteins, the cognate YAP/TAZ-binding transcription factors, could bind to KIRREL1 gene (Fig. 6i). Thus, KIRREL1 is not only an upstream regulator of the Hippo pathway, but also a target gene of YAP/TAZ and constitutes a negative feedback loop to control Hippo signaling. Activation of YAP/TAZ induces the transcription of *KIRREL1* gene, and KIRREL1 localizes to tight junctions where KIRREL1 coordinates LATS1/2 phosphorylation by MST1/2, thus preventing uncontrolled YAP/TAZ activation.

**Fig 6.**
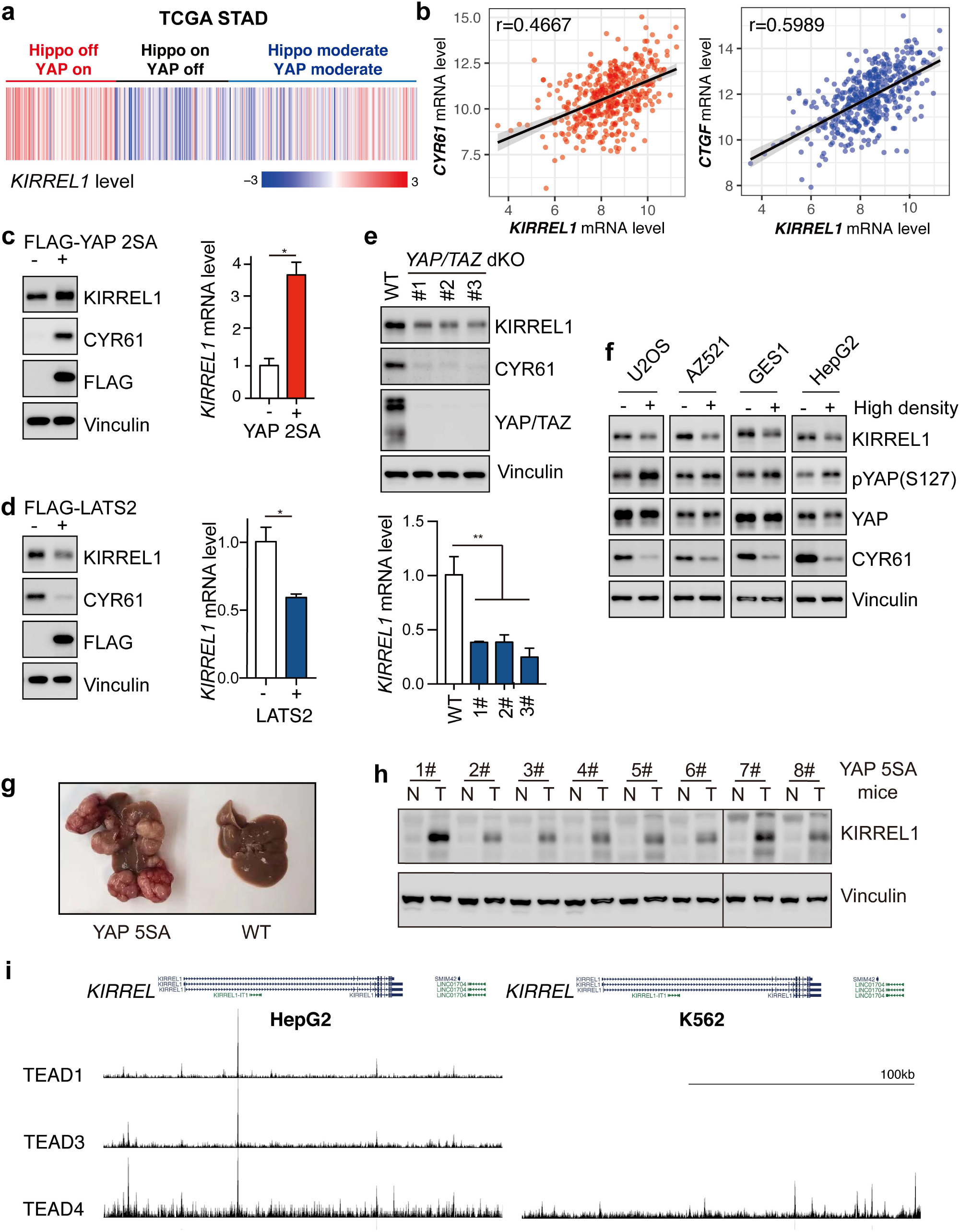
*KIRREL1* is a YAP/TAZ target gene. (A) KIRREL1 mRNA level in Hippo off/YAP on, Hippo on/YAP off, and Hippo moderate/YAP moderate stomach cancer patients defined in Supplementary Data Fig. S1e. (B) The correlation between KIRREL1 mRNA level and *CYR61* (left) and *CTGF* (right) mRNA levels in TCGA stomach cancer samples. (C) Overexpression of YAP 2SA increased KIRREL1 protein (left) and mRNA (right) levels in HGC-27 cells. (D) Overexpression of LATS2 decreased KIRREL1 protein (left) and mRNA (right) levels in HGC-27 cells. (E) YAP/TAZ double KO significantly decreased KIRREL1 protein (left) and mRNA (right) levels in HGC-27 cells. (F) Hippo activation by high cell density led to decreased KIRREL1 level in U2OS, AZ521, GES1, and HepG2 cells. (G) IB showed KIRREL1 level significantly increased in liver cancer tissues induced by YAP 5SA hydrodynamic injection. (H) Representative image of YAP 5SA mice livers. (I) Chip-seq signals of TEAD in HepG2 and K562 cell lines. Mean and standard error were presented (*p<0.05, **p < 0.01, ***p < 0.001, ****p <0.0001, ns = not significant; n=2; t test).

### p35-KIRREL1 is a regulator of Hippo signaling and is sensitively targeted by YAP/TAZ

Immunoblotting showed that a ~35 kDa protein band was frequently observed when KIRREL1 protein levels were examined in different cells. The antibody used to detect KIRREL1 recognizes the C-terminal sequences of KIRREL1, and this protein band completely disappeared in KIRREL1 knockout cells, indicating that this smaller protein would be a C-terminal fragment of KIRREL1. We dubbed this protein as p35-KIRREL1 (labeled as P35 in figures) (Fig. 7a,b). Compared to long KIRREL1 isoforms, p35-KIRREL1 was much more dynamically regulated by YAP/TAZ activity, in a manner comparable to CYR61 (Fig. 7c-f). The expression of p35-KIRREL1 was robustly reduced when cells were cultured at high density, and nearly disappeared in *YAP/TAZ* or *TEAD1-4* knockout cells (Fig. 7c,d). In contrast, p35-KIRREL1 protein levels were significantly elevated in cells expressing YAP, especially active YAP (YAP-2SA) (Fig. 7e,f). We also noticed that when KIRREL-i1 was ectopically expressed, it produced a smaller protein band identical to p35-KIRREL1 (Fig. 7g). We deleted inframe ATG codons in the C-terminal region of *KIRREL1* and observed that only the 1384-1386bp ATG (encode methionine 462, M462) deletion mutant completely blocked the generation of the small KIRREL1 protein (Fig, 7a,g), suggesting that p35-KIRREL1 was likely an alternatively translated protein including C-terminal 296aa of KIRREL1 (start from M462).

**Fig 7.**
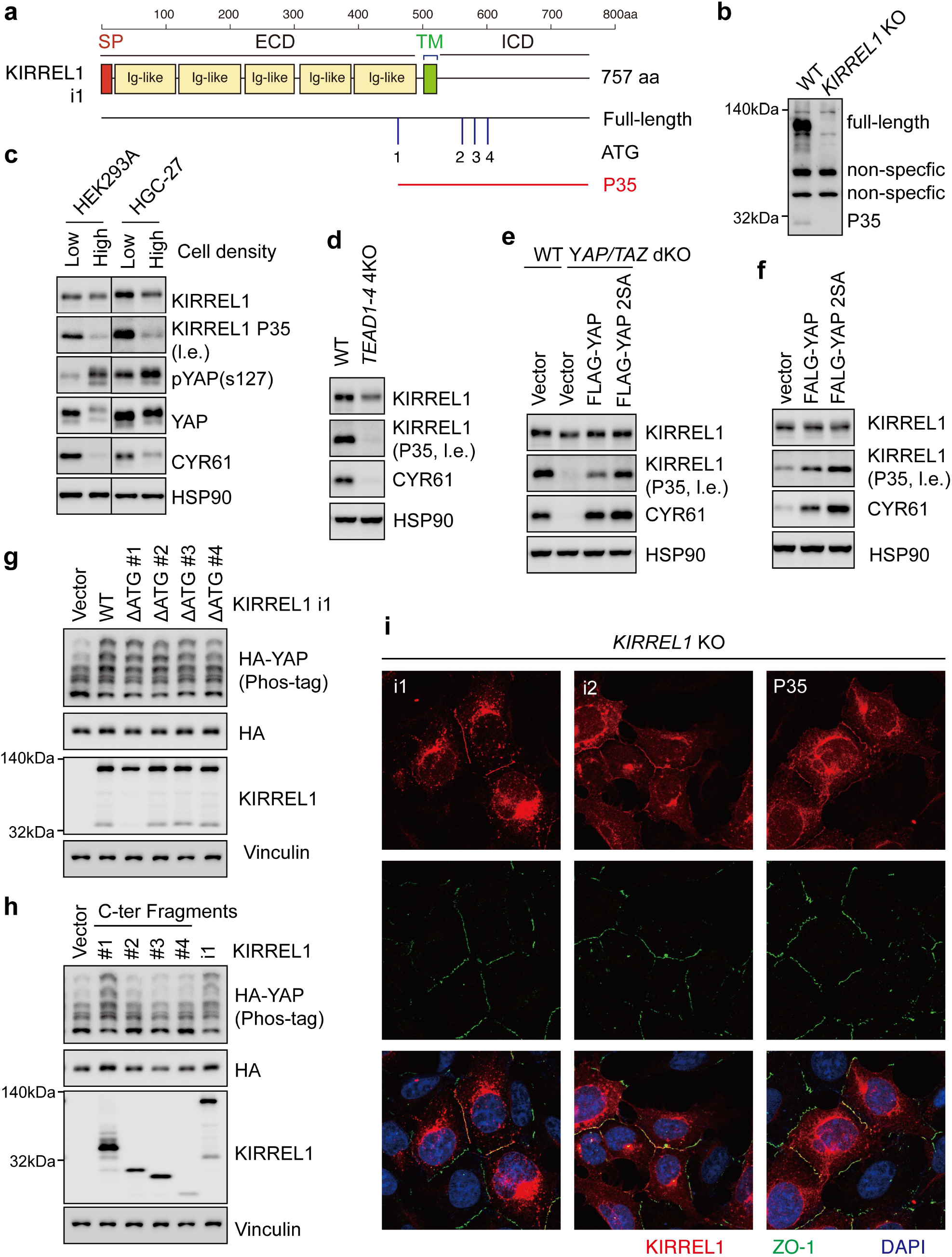
p35-KIRREL1 is a regulator of Hippo signaling and is sensitively targeted by YAP/TAZ. (A) Alignment of KIRREL1 P35 and different ΔATG mutants in full-length KIRREL1. (B) IB of KIRREL1 in wild-type and KIRREL1 KO HEK293A cells. Bands shared by two lanes would be nonspecific IB signals. KIRREL1 P35 was shown as a ~35kD band. (C) Hippo activation by high cell density significantly decreased KIRREL1 P35 level in HEK293A and HGC-27 cells. l.e. indicated longer exposure time. (D) TEAD1-4 knock out significantly decreased KIRREL1 P35 level in HEK293A cells. (E) YAP/TAZ knock out significantly decreased KIRREL1 P35 level, while YAP and YAP 2SA rescue increased KIRREL1 P35 level in HEK293A cells. (F) YAP and YAP 2SA overexpression significantly increased KIRREL1 P35 level in HEK293A cells. (G) Overexpression of different ΔATG KIRREL1 isoform 1 mutants in KIRREL1 KO HEK293A cells and their effects on YAP phosphorylation. ΔATG1: Δ1384-1386 bp; ΔATG2: Δ1648-1650 bp; ΔATG3: Δ1702-1704 bp; ΔATG4: Δ1792-1794 bp. ΔATG1 KIRREL1 mutant lost P35 band of KIRREL1. (H) Overexpression of different P35 sequences corresponding to 4 candidate ATG sites and their effects on YAP phosphorylation. P35 1# inhibited YAP like full-length isoform 1 did. (I) KIRREL1 P35 showed similar localization in tight junction with full-length isoform 1 and isoform 2. No-tag KIRREL1 P35, full-length isoform 1 and isoform 2 were expressed in KIRREL1 KO HEK293A cells, and cells were fixed for immunofluorescence staining.

As shown earlier, KIRREL1 gene has at least four isoforms. By analyzing the ENCODE ChIP-Seqdata, we observed that in HepG2 cells, TEAD1/2/4 bound to multiple regions of *KIRREL1* gene, with a dominant peak localized inside the first intron, which likely controls the expression of *KIRREL1-i1/2/3,* and another peak in the 3’ region, which may mostly control the expression of *KIRREL1-i4* (Fig. 6i). Moreover, in K562 cells, TEAD4 ChIP-seq signals were enriched in the 3’ region (Fig. 6i). In principle, all isoforms of KIRREL1 containing TM domain and ICD should be able to produce p35-KIRREL1 (Fig. 7a). We propose that transcription of different *KIRREL1* isoforms might be induced by YAP/TAZ, and shorter isoforms, especially isoform 4, may predominantly translate p35-KIRREL1. In this way, the effect of YAP/TAZ on p35-KIRREL1 protein could be amplified compared to long isoforms.

We then asked if p35-KIRREL1 played a role in regulating the Hippo pathway. We expressed p35-KIRREL1 in *KIRREL1* knockout cells, and observed robust induction of YAP phosphorylation. However, three other genes encoding different ICD regions of KIRREL1 (by alternative translation start codon) failed to boost YAP phosphorylation (Fig. 7a,h). The TM sequence in p35-KIRREL1 was intact, and p35-KIRREL1 was localized at tight junctions, similar to KIRREL1-i1/2 (Fig. 7i). Thus, the ICD domain of KIRREL1 is not sufficient to regulate the Hippo pathway, and the localization of KIRREL1 is strictly required.

### KIRREL1 functions as a tumor suppressor and inhibits liver tumorigenesis

Next, we evaluated the effect of KIRREL1 on the proliferation of cancer cells. As a negative regulator of YAP, *KIRREL1* knockout should promote the proliferation of cancer cells after YAP is activated. To gain a comprehensive view of the role of KIRREL1 in cell proliferation, we analyzed datasets from the Cancer Dependency Map project (http://depmap.org). This project used genome-wide RNAi and CRISPR/Cas9 loss-of-function screens to systematically identify essential genes across hundreds of human cancers. The effect of single gene knockout on cell survival and proliferation was calculated to yield a dependency score for each cell line, in which a positive score indicated that a gene knockout led to increased cell proliferation. Genetic co-dependency is identified if some genes are commonly required by a set of cancer cell lines. Strikingly, we found that the Hippo pathway genes *NF2, AMOTL2* and *LATS2* were the top 3 candidates showing high co-dependencies with *KIRREL1* (Fig. 8a). Moreover, *KIRREL1* was identified as the 2^nd^ top candidate with high co-dependencies with *LATS2* and *SAV1* (Fig. 8a). These data indicate that KIRREL1 and NF2, AMOTL2, LATS2, and SAV1 have similar inhibitory effects on cell proliferation and may very likely function in a same pathway.

**Fig 8.**
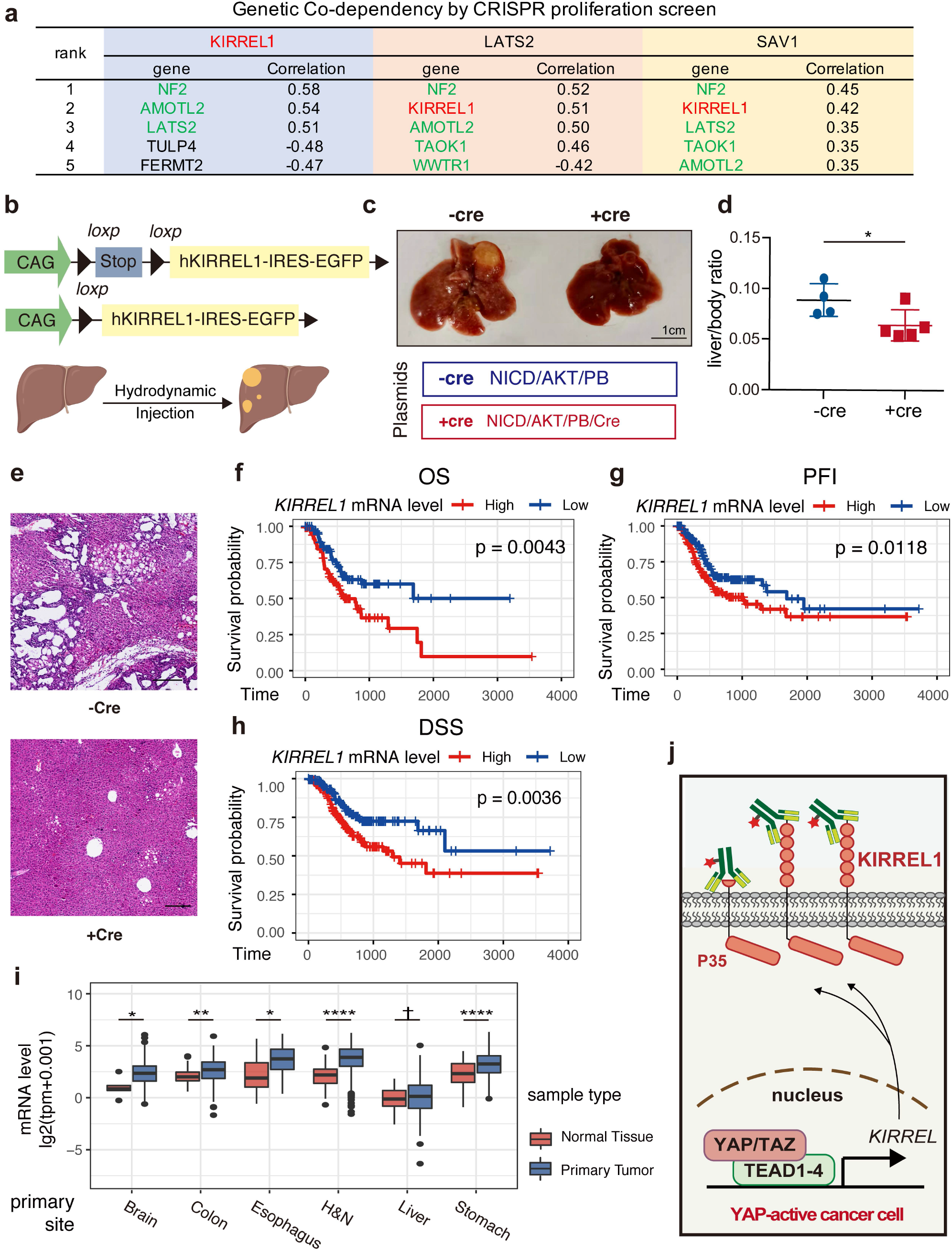
KIRREL1 inhibits tumorigenesis and represents a potential therapeutic target. (A) Top 5 genes sharing genetic co-dependency with KIRREL1, LATS2, and SAV1 are listed. KIRREL1 marked in red. Known Hippo pathway regulators marked in green. Correlations were calculated by Pearson’s correlation of two gene’s dependency scores in more than 1000 cell lines from Depmap.org datasets. (B) Schemes of R26-e(CAG-LSL-hKIRREL1 isoform2-IRES-EGFP-PolyA) mouse model and mouse cholangiocarcinoma model induced by hydrodynamic injection of NICD, AKT, and piggyBac (PB) via lateral tail vein of mice. Cre plasmids were also injected in +Cre groups to induce the expression of hKIRREL1 isoform2. Tissue samples were harvested 10 weeks after injection. (C) Representative image of -/+Cre mice livers. While hydrodynamic injection led to yellow blister lesions in liver, coinjection of Cre significantly relieved lesions. (D) Liver/body weight ratio of -/+Cre mice livers (*p <0.05, t test). (E) Representative image of H&E staining in -/+Cre mice livers. (F) Prognostic value of *KIRREL1* level in stomach cancer. TCGA stomach cancer patients were divided equally into high/low *KIRREL1* expression groups. Their overall survival (OS) was compared by log rank test. (G) Prognostic value of *KIRREL1* level in stomach cancer. TCGA stomach cancer patients were divided equally into high/low *KIRREL1* expression groups. Their progression-free interval (PFI) was compared by log rank test. (H) Prognostic value of *KIRREL1* level in stomach cancer. TCGA stomach cancer patients were divided equally into high/low *KIRREL1* expression groups. Their disease-specific survival (DSS) was compared by log rank test. (I) *KIRREL1* level and its difference between tumor and normal tissues in different cancer types. Data from TCGA pan-cancer datasets. (J) Model of KIRREL1 as a therapeutic antibody target in YAP-active cancers.

To further explore whether of KIRREL1 could inhibit tumorigenesis *in vivo,* we generated a transgenic mouse line that conditionally expresses *KIRREL1*-i2 (Fig. 8b). An intrahepatic cholangiocarcinoma (iCCA) model was then established by hydrodynamic injection of *piggyBac* (PB) plasmids expressing cancer drivers, including a truncated Notch receptor (NICD) and myristoylated AKT (myr-AKT), with or without a Cre-expressing plasmid, into livers of *KIRREL1* transgenic mice (Fig. 8b). While expression of NICD and myr-AKT effectively induced iCCA, as indicated by the presence of extensive bile duct structures, co-injection of *Cre* plasmids, which turned on *KIRREL1* expression, blocked the formation of iCCA (Fig. 8c-e). Thus, the expression of KIRREL1 can effectively inhibit tumorigenesis.

As a direct target of YAP/TAZ, the expression of KIRREL1 should increase in YAP/TAZ-active tumors. Indeed, KIRREL1 was significantly upregulated in STAD tumor samples, and patients with high *KIRREL1* expression have shorter overall survival (OS), progression-free intervals (PFI), and disease-specific survival (DSS) (Fig. 8f-h). Moreover, *KIRREL1* mRNA level was significantly elevated in tumor samples compared to normal tissues in different cancer types, including brain, colon, esophagus, and others (Fig. 8j).

## Discussion

To this date, the best known function of KIRREL1 is the regulation of glomerular permeability via its interaction with nephrins^31^. Our work has added new insights into the physiological and pathological functions of KIRREL1 and demonstrated that this protein is an important regulator of the Hippo pathway. Localized at tight junctions, KIRREL1 interacts with key Hippo pathway components SAV1 and LATS1/2 via its C-terminal domains, promoting the simultaneous recruitment of LATS1/2 and SAV1 to the plasma membrane where LATS1/2 are activated by MST1/2 (Fig. 3). KIRREL1 is active only when it is localized to tight junctions and can thus work as a cell density sensor and a Hippo signaling switch (Figs. 4,5). Moreover, *KIRREL1* is also a target gene of YAP. YAP induces the expression of *KIRREL1*, in turn activating Hippo signaling and inhibiting YAP activity, and thus forming a negative feedback loop to maintain homeostasis of Hippo signaling (Fig. 6). The expression of *KIRREL1* is limited to specific cell types, including endometrial stromal cells, type 1 alveolar cells, smooth muscle cells, fibroblasts, breast myoepithelial cells, and adipocytes (single cell sequencing data, https://www.proteinatlas.org). Hence, the role of KIRREL1 in the regulation of the Hippo pathway should be tissue-specific, especially at physiological conditions, and this warrants future studies.

By inhibiting YAP/TAZ activity, KIRREL1 represents a tumor suppressor. Indeed, downregulation of KIRREL1 in multiple cancer cells induced cell proliferation, as shown in an unbiased screen in Cancer Dependency Map project (http://depmap.org). Likewise, as demonstrated in this study, transgenic expression of *KIRREL1* effectively blocked tumorigenesis in a mouse iCCA model (Fig. 8). Hence, induction of *KIRREL1* expression represents a novel strategy to treat different types of cancers. In this study, we have identified a p35-KIRREL1 protein that can also inhibit YAP/TAZ activity and is relatively small in size (Fig. 7). It would be interesting to explore the anti-cancer potential of p35-KIRREL1 protein in future studies.

The Hippo signaling pathway is an attractive target for cancer therapy. However, one challenge in this field is the lack of a well-established cell surface marker or regulator for therapeutic targeting ^18^. As a plasma membrane protein, KIRREL1 can serve as a cell surface marker of YAP-active tumor cells. In particular and as shown in this study, p35-KIRREL1 is highly sensitive to YAP activity. The development of specific antibodies against ECD of KIRREL1 isoforms and p35-KIRREL1 can help advance our understanding of the relationship between KIRREL1 and YAP/TAZ in cancer. These antibodies may be used directly to modulate Hippo pathway strength by targeting KIRREL1, or as antibody-drug conjugates (ADCs) to deliver cancer drugs specifically to tumors (Fig. 8j). In summary, our work established KIRREL1 as a cell surface regulator of the Hippo pathway and a potential biomarker and drug target in YAP-driven tumors.

Authors note: while this manuscript is in preparation, a study with partially overlapping findings has been published^32^.

## Author Contributions

Y.G., and F.X.Y. designed the experiments and wrote the manuscript. Y.G., Y.W., Z.S., J.L., performed experiments and analyzed data. C.H., and F.L., analyzed of ChIP-seq data. F.X.Y. supervised this study.

## Acknowledgments

This study is supported by grants from the Ministry of Science and Technology of China (National Key R&D program, 2018YFA0800304, and 2020YFA0803202), the National Natural Science Foundation of China (81772965), and Science and Technology Commission of Shanghai Municipality (19JC1411100, 21S11905000) to F.-X.Y.

## Methods

### Mouse Models

All mouse experiments were approved by the Animal Ethics Committee of Shanghai Medical College, Fudan University and carried out in accordance with institutional guidelines. R26-e(CAG-LSL-hKIRREL1 isoform2-IRES-EGFP-PolyA) C57BL/6 mice were in-house generated as described in Fig. 8b. Liver cancer models were established by hydrodynamic injection of plasmids expressing cancer drivers or transposase. In detail, 25 μg NICD, 25 μg myristoylated AKT, 10 μg piggyBac (PB), and 100 μg Cre (or control) plasmids were diluted in 2 ml 0.9% NaCl, filtered, and hydrodynamically injected via lateral tail vein with the final volume as 10% (in ml) of the total body weight (in grams). Retro-orbital blood was collected and serum was prepared for blood chemistry. In experiments involving tumorigenesis, mice were euthanized when severe abdominal enlargement or body weight loss (70% of control littermates) was observed. At the indicated age, mouse livers harvested were either fixed in formalin solution, paraffin embedded, and sectioned for pathological analysis, or stored in liquid nitrogen and used for gene expression analysis.

### Cell lines and cell culture

HEK293A and HepG2 cells were maintained in DMEM medium supplemented with 10% fetal bovine serum (FBS). HGC-27, U2OS, AZ521, and GES1 cells were maintained in RPMI 1640 medium supplemented with 10% FBS. All culture medium was supplemented with 50 mg/ml penicillin/streptomycin. Cells were incubated at 37°C with 5% CO2. NF2 KO, SAV1 KO, MST1/2 dKO, LATS1/2 dKO, MOB1A/B dKO, WWC1/2/3 tKO, and MAP4K3-7 5KO HEK293A cells were described previously^33,34^. KIRREL KO HEK293A cells were generated in this study using CRISPR/Cas9 system. The deletion of genes in each cell line was confirmed by immunoblotting or DNA sequencing. Cells stably expressing different target genes were established by lentiviral transduction.

### CRISPR/Cas9-mediated gene editing

pSpCas9(BB)-2A-Puro (PX459) V2.0 (Addgene #62988) was provided by Feng Zhang ^35^. Genespecific sgRNAs were designed using the CRISPR design tool at http://www.genome-engineering.org/crispr. HEK293A cells were transfected and selected with puromycin for 2–3 days, and single cell was seeded into 96-well plates. Monoclonal cells were screened by immunoblotting and/or Sanger sequencing (Extended Data Figure 8). The single-guide RNA (sgRNA) sequences were listed in Table S5.

### Plasmids, transfection, and lentivirus production

Genes in the Hippo pathway were subcloned into pLVX-puro vector (Takara, #632164) with HA-, MYC- or FLAG-tag, and used for transient overexpression. Transfection was performed using PolyJet™ DNA In Vitro Transfection Reagent (Signagen, # SL100688) following manufacturer’s instructions. For lentivirus production, pLVX-based plasmids were co-transfected into HEK293A cells together with PsPAX2 and pMD.2g. Two days later, medium containing virus was filtered and used for viral transduction.

### BioID-MS

HEK293A cells stably expressing BirA-tag Hippo regulators were incubated with complete medium containing 50 uM Biotin for 12 hours before lysis, sonication, and centrifugation. Equilibrated streptavidin dynabeads were mixed with supernatant of cell lysates and incubated overnight. Beads were then washed 3 times with lysis buffer before 1×SDS loading buffer was added to elute the proteins. Target proteins were separated by sodium dodecyl sulfate–polyacrylamide gel electrophoresis (SDS-PAGE). Gels were cut and sent for mass spectrometry analysis.

### Immunoblotting

Whole cell lysates (~10 μg proteins/sample) were separated by SDS-PAGE, and proteins were transferred onto nitrocellulose membranes. Following blocking in 5% non-fat milk, membranes were incubated with primary antibodies in 5% bovine serum albumin (BSA) overnight at 4°C, and then with secondary antibodies in 5% milk for 1 h at room temperature. High-signal ECL Western Blotting Substrate (Tanon, #180-501) was applied onto the membranes, and chemiluminescence was detected using a Tanon 5200S imaging system. NEPH1 antibody (F-6, sc-373787, SCBT) was used in immunoblotting and immunofluorescence assays to detect KIRREL.

### Immunoprecipitation

Whole cell lysates were prepared in mild lysis buffer (50 mM HEPES at pH 7.5, 150 mM NaCl, 1 mM EDTA, 1% NP-40, 10 mM pyrophosphate, 10 mM glycerophosphate, 50 mM NaF, and 1.5 mM Na3VO4) supplemented with 1 mM PMSF and protease and phosphatase inhibitor cocktails (Biotool), then incubated with FLAG- or HA-tag antibodies for 1 hour at 4°C. Protein A agarose beads (Repligen, 10-1003-02) or protein A/G magnetic beads (Pierce, 88803) were added and incubated for another 1 h to capture protein-antibody complex. Beads were washed 4 times with mild lysis buffer, and precipitated proteins were dissolved in 1× SDS-PAGE sample buffer and analyzed by immunoblotting.

### Immunofluorescence

Cells grown on cover slides were fixed in PBS containing 4% formaldehyde for 10 min and then treated with 0.1% Triton X-100 for 15 min. After blocking with 3% goat serum, cells were stained with primary antibodies in PBST with 3% BSA overnight at 4°C, and secondary antibodies at room temperature for 1 h. Cover slides were then mounted with ProLong™ Gold Antifade Mountant with DAPI (ThermoFisher, P36935). Images were captured with Leica SP8 or Zeiss LSM900 confocal microscope.

### Purification of recombinant proteins and kinase assay

GST-YAP was expressed in *Escherichia coli* (BL-21). Recombinant proteins were purified by affinity chromatography using glutathione (GSH) agarose beads (GE Healthcare) and dialyzed in mild lysis buffer. Purified proteins were snap frozen in liquid nitrogen and stored at −80°C in aliquots.

HA-LATS1/2 was immunoprecipitated, and proteins on beads were washed twice with the kinase assay buffer (30 mM HEPES, 50 mM potassium acetate, 5 mM MgCl2). GST-YAP (0.5 μg/reaction) was used as substrate for LATS1. The kinase reaction was carried out in the presence of 500 μM cold ATP for 30 min at 30°C, then terminated with SDS sample buffer, and subjected to Western blotting. Phospho-specific antibody for pYAP-S127 was used to evaluate the kinase activity of LATS1.

### RNA extraction, reverse transcription, and real-time PCR

Total RNA was extracted using the RNeasy Plus mini kit (Qiagen). cDNA was generated by the PrimeScript RT reagent Kit (TaKaRa), and quantitative qPCR was conducted using SYBR Green qPCR Master Mix (TaKaRa) on the 7500 Real-Time PCR system (Applied Biosystems). Primers used in PCR were as follows: KIRREL, F:5’-ATGGAGGCCGACTTTCAGAC-3’, R:5’-GATGGTGGCCCCAGCTATGA-3’; CYR61, F:5’-AGCCTCGCATCCTATACAACC-3’, R:5’-TTCTTTCACAAGGCGGCACTC-3’; CTGF, F:5’-CCAATGACAACGCCTCCTG-3’, R:5’-TGGTGCAGCCAGAAAGCTC-3’; ANKRD1, F:5’-CACTTCTAGCCCACCCTGTGA-3’, R:5’-CCACAGGTTCCGTAATGATTT-3’; AMOTL2, F:5’-AGCTTCAATGAGGGTCTGCT-3’, R:5’-TGAAGGACCTTGATCACTGC −3’; GAPDH, F:5’-ATGGGGAAGGTGAAGGTCG-3’, R:5’-GGGGTCATTGATGGCAACAATA −3’.

### Data collection and survival analysis

Processed TCGA pan-cancer, stomach cancer, GTEx normal tissue RNAseq data, clinical data, and somatic mutation datasets were downloaded from UCSC Xena datasets^36^. Raw data are available at GDC Data Portal (https://portal.gdc.cancer.gov)^37^ and GTEx portal (https://www.gtexportal.org/). Survival data were analyzed by Kaplan-Meier analysis and Cox proportional hazard analysis. Analysis and visualization (R package ggplot2) were done by R (version 3.6.2).

### Statistical analysis

Statistical analyses were performed using GraphPad Prism 8 software (GraphPad Software, Inc, USA) or R. Experiments were repeated for at least two times as reported in the figures and corresponding figure legends. The results were expressed as the mean ± SEM. Statistical significance was determined using Student’s *t*-test between groups. *p < 0.05, **p < 0.01, ***p < 0.001, ****p < 0.0001, n.s. indicates not significant.

### Data availability

All unique/stable reagents generated in this study are available from the lead contact with a completed Uniform Biological Materials Transfer Agreement. This paper analyzes existing, publicly available data. Processed TCGA pan-cancer, stomach cancer, GTEx normal tissue RNAseq data, clinical data and somatic mutation datasets were downloaded from UCSC Xena datasets. Raw data was available from GDC Data Portal (https://portal.gdc.cancer.gov) and GTEx portal.

---

**Fig. S1.**
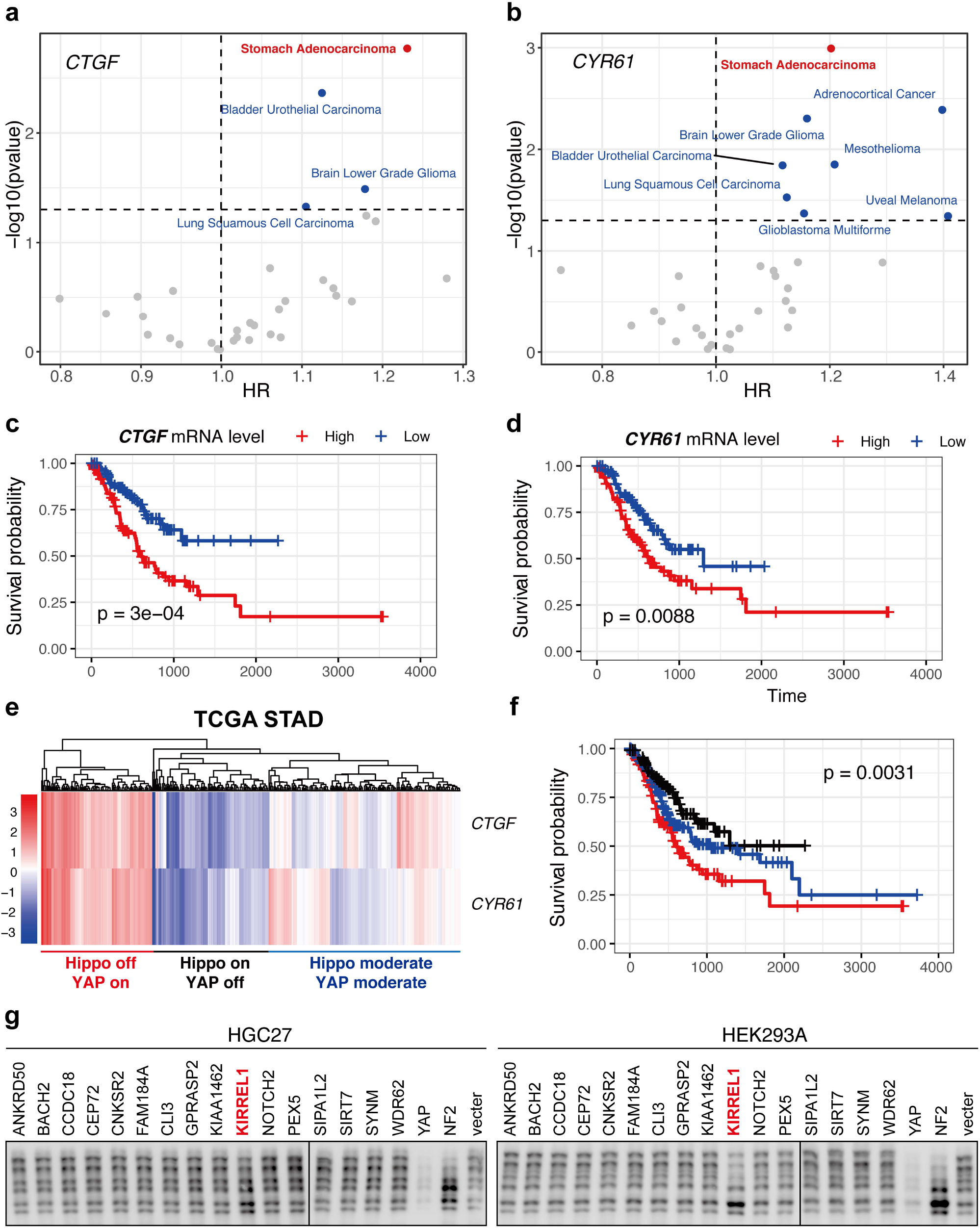
Prognostic value of the Hippo pathway in different cancer types, related to Fig. 1. (A) Prognostic value of *CTGF* level in different cancer types. Hazard ratio (HR) and its *p* value was calculated by Cox proportional hazard regression model based on TCGA pan-cancer datasets. (B) Prognostic value of *CYR61* level in different cancer types. (C) Prognostic value of *CTGF* level in stomach cancer. TCGA stomach cancer patients were divided equally into high/low *CTGF* expression groups. Their survival was compared by log rank test. (D) Prognostic value of *CYR61* level in stomach cancer. (E) TCGA stomach cancer patients were classified into Hippo off/YAP on, Hippo on/YAP off, and Hippo moderate/YAP moderate groups by an enrichment analysis of *CTGF* and *CYR61* levels. (F) Hippo off/YAP on stomach cancer patients had a worse prognosis. Survival of three groups of patients was compared by log rank test. (G) YAP Phos-tag blotting result of secondary genetic screen.

**Fig. S2.**
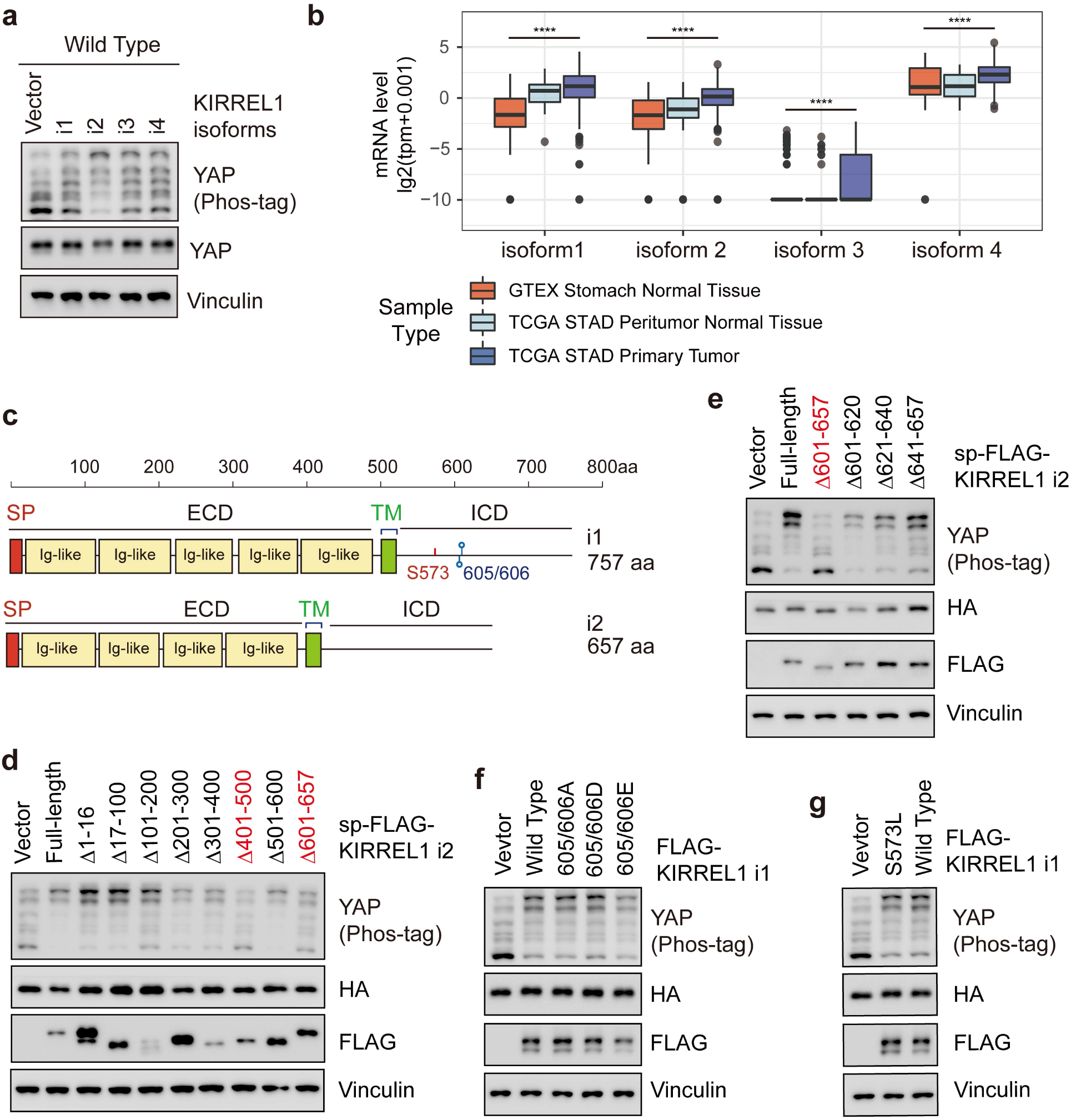
Mapping the key domains in KIRREL1 that are important for the regulation of YAP, related to Fig.2. (A) All isoforms of KIRREL1 increased YAP phosphorylation level, while isoform 2 showed the strongest effect in wild-type HEK293A cells. (B) mRNA levels of different KIRREL1 isoforms in TCGA peritumor normal tissue, stomach cancer, and GTEX stomach normal tissue (****p <0.0001, t test). (C) Structure of KIRREL i1 and i2. (D) Overexpression of KIRREL1 mutants each deleting 100 aa in KIRREL1 KO HEK293A cells showed their effects on YAP phosphorylation. sp-FLAG-KIRREL1 indicated knocking in a FLAG tag sequence after the sequence of 1-16aa predicted signal peptide in KIRREL1 plasmid. Mutants losing function marked in red. (E) Overexpression of KIRREL1 mutants each deleting 20aa among 601-657aa in KIRREL1 KO HEK293A cells showed all mutants had effects on YAP phosphorylation. (F) Overexpression of KIRREL1 mutants with FYN phosphorylation sites mutated in KIRREL1 KO HEK293A cells could still increase YAP phosphorylation. (G) Overexpression of KIRREL1 S537 mutant in KIRREL1 KO HEK293A cells could still increase YAP phosphorylation.

**Fig. S3.**
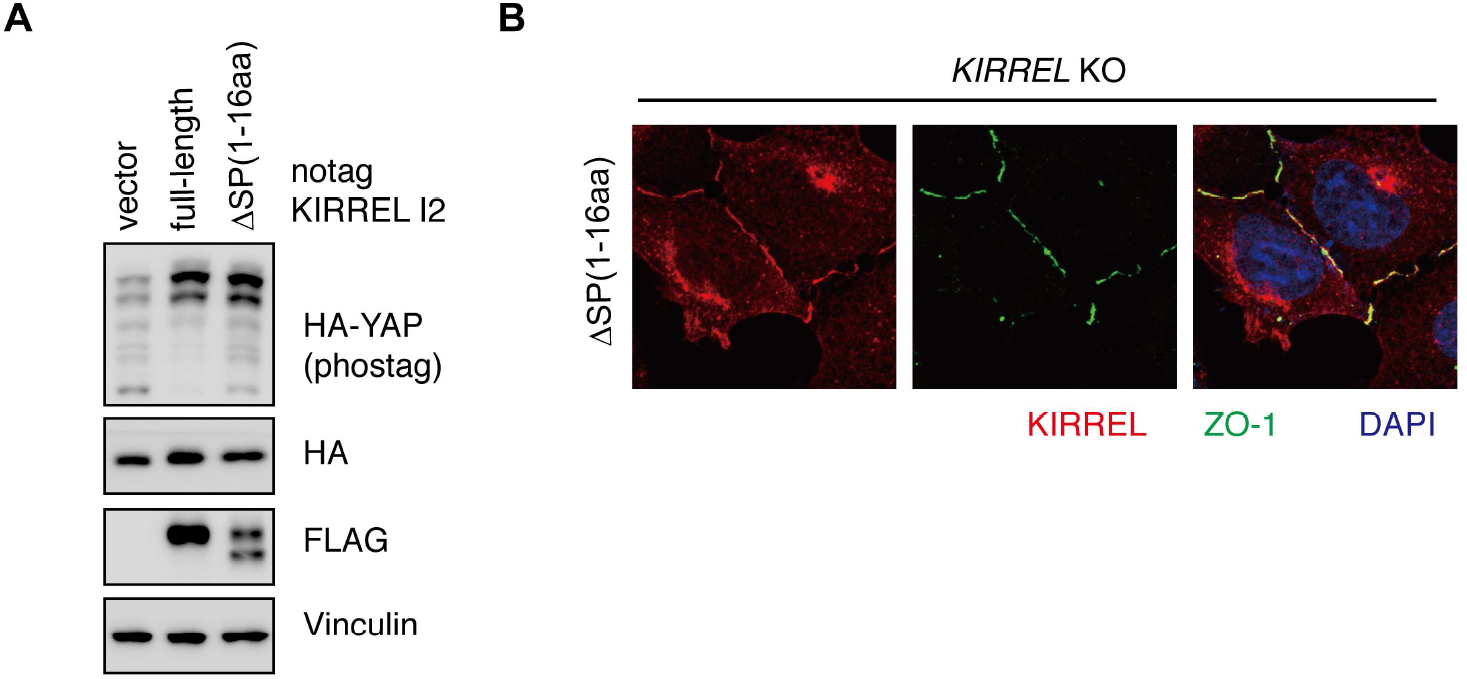
SP domain is dispensable for KIRREL1’s localization and function, related to Fig.4. (A) Overexpression of KIRREL mutants deleting 401-420aa showed SP domain was dispensable for KIRREL1 function. (B) Immunofluorescence staining in *KIRREL1* KO HEK293A cells showed SP-domain-deleted mutant lost tight junction co-localization.

## References

1. Driskill, J.H. & Pan, D. The Hippo Pathway in Liver Homeostasis and Pathophysiology. Annu Rev Pathol 16, 299–322 (2021).

2. Russell, J.O. & Camargo, F.D. Hippo signalling in the liver: role in development, regeneration and disease. Nat Rev Gastroenterol Hepatol (2022).

3. Yu, F.X., Zhao, B. & Guan, K.L. Hippo Pathway in Organ Size Control, Tissue Homeostasis, and Cancer. Cell 163, 811–28 (2015).

4. Zheng, Y. & Pan, D. The Hippo Signaling Pathway in Development and Disease. Dev Cell 50, 264–282 (2019).

5. Wang, Y., Yu, A. & Yu, F.X. The Hippo pathway in tissue homeostasis and regeneration. Protein Cell, 349–359 (2017).

6. Camargo, F.D. et al. YAP1 increases organ size and expands undifferentiated progenitor cells. Curr Biol 17, 2054–60 (2007).

7. Zhou, D. et al. Mstl and Mst2 maintain hepatocyte quiescence and suppress hepatocellular carcinoma development through inactivation of the Yapl oncogene. Cancer Cell 16, 425–38 (2009).

8. Heallen, T. et al. Hippo pathway inhibits Wnt signaling to restrain cardiomyocyte proliferation and heart size. Science 332, 458–61 (2011).

9. Nishio, M. et al. Cancer susceptibility and embryonic lethality in Mobla/lb double-mutant mice. J Clin Invest 122, 4505–18 (2012).

10. Yu, F.X. et al. Regulation of the Hippo-YAP pathway by G-protein-coupled receptor signaling. Cell 150, 780–91 (2012).

11. Chan, S.W. et al. A role for TAZ in migration, invasion, and tumorigenesis of breast cancer cells. Cancer Res 68, 2592–8 (2008).

12. Mo, J.S. et al. Cellular energy stress induces AMPK-mediated regulation of YAP and the Hippo pathway. Nat Cell Biol 11, 500–10 (2015).

13. Li, H. et al. *YAP/TAZ.* drives cell proliferation and tumour growth via a polyamine-elF5A hypusination-LSDl axis. Nat Cell Biol (2022).

14. Pearson, J.D. et al. Binary pan-cancer classes with distinct vulnerabilities defined by pro- or anti-cancer YAP/TEAD activity. Cancer Cell 39, 1115–1134 e12 (2021).

15. Zanconato, F., Cordenonsi, M. & Piccolo, S. YAP/FAZ at the Roots of Cancer. Cancer Cell 29, 783–803 (2016).

16. Jiao, S. et al. Targeting IRF3 as a YAP agonist therapy against gastric cancer. J Exp Med 215, 699–718 (2018).

17. Dong, J. et al. Elucidation of a universal size-control mechanism in Drosophila and mammals. Cell 130, 1120–33 (2007).

18. Johnson, R. & Halder, G. The two faces of Hippo: targeting the Hippo pathway for regenerative medicine and cancer treatment. Nat Rev Drug Discov 13, 63–79 (2014).

19. Liu-Chittenden, Y. et al. Genetic and pharmacological disruption of the TEAD-YAP complex suppresses the oncogenic activity of YAP. Genes Dev 26, 1300–5 (2012).

20. Tang, T.T. et al. Small Molecule Inhibitors of TEAD Auto-palmitoylation Selectively Inhibit Proliferation and Tumor Growth of NF2-deficient Mesothelioma. Mol Cancer Ther (2021).

21. Lin, C.Y. et al. Membrane protein-regulated networks across human cancers. Nat Commun 10, 3131 (2019).

22. Zhao, B. et al. Inactivation of YAP oncoprotein by the Hippo pathway is involved in cell contact inhibition and tissue growth control. Genes Dev 21, 2747–61 (2007).

23. Donoviel, D.B. et al. Proteinuria and perinatal lethality in mice lacking NEPH1, a novel protein with homology to NEPHRIN. Mol Cell Biol 21, 4829–36 (2001).

24. Harita, Y. et al. Neph1, a component of the kidney slit diaphragm, is tyrosine-phosphorylated by the Src family tyrosine kinase and modulates intracellular signaling by binding to Grb2. J Biol Chem 283, 9177–86 (2008).

25. Solanki, A.K. et al. Mutations in KIRREL1, a slit diaphragm component, cause steroid-resistant nephrotic syndrome. Kidney Int 96, 883–889 (2019).

26. Callus, B.A., Verhagen, A.M. & Vaux, D.L. Association of mammalian sterile twenty kinases, Mstl and Mst2, with hSalvador via C-terminal coiled-coil domains, leads to its stabilization and phosphorylation. FEBS J 273, 4264–76 (2006).

27. Yin, F. et al. Spatial organization of Hippo signaling at the plasma membrane mediated by the tumor suppressor Merlin/NF2. Cell 154, 1342–55 (2013).

28. Wang, Y. et al. Stabilization of Motin family proteins in NF2-deficient cells prevents full activation of YAP/TAZ and rapid tumorigenesis. Cell Rep 36, 109596 (2021).

29. Zhao, B., Li, L., Tumaneng, K., Wang, C.Y. & Guan, K.L. A coordinated phosphorylation by Lats and CK1 regulates YAP stability through SCF(beta-TRCP). Genes Dev 20, 72–85 (2010).

30. Guo, X. et al. Single tumor-initiating cells evade immune clearance by recruiting type II macrophages. Genes Dev 31, 247–259 (2017).

31. Liu, G. et al. Neph1 and nephrin interaction in the slit diaphragm is an important determinant of glomerular permeability. J Clin Invest 112, 209–21 (2003).

32. Paul, A. et al. Cell adhesion molecule KIRREL1 is a feedback regulator of Hippo signaling recruiting SAV1 to cell-cell contact sites. Nat Commun 13, 930 (2022).

33. Meng, Z. et al. MAP4K family kinases act in parallel to MST1/2 to activate LATS1/2 in the Hippo pathway. Nat Commun 6, 8357 (2015).

34. Plouffe, S.W. et al. Characterization of Hippo Pathway Components by Gene Inactivation. Mol Cell 64, 993–1008 (2016).

35. Ran, F.A. et al. Genome engineering using the CRISPR-Cas9 system. Nat Protoc 3, 2281–2308 (2013).

36. Goldman, M.J. et al. Visualizing and interpreting cancer genomics data via the Xena platform. Nat Biotechnol 38, 675–678 (2020).

37. Hmeljak, J. et al. Integrative Molecular Characterization of Malignant Pleural Mesothelioma. CancerDiscov 8, 1548–1565 (2018).

